# Efficient generation of mNeonGreen *Plasmodium falciparum* reporter lines enables quantitative fitness analysis

**DOI:** 10.1101/2022.07.11.499328

**Authors:** Johanna Hoshizaki, Hannah Jagoe, Marcus Lee

## Abstract

CRISPR editing has enabled the rapid creation of fluorescent *Plasmodium* transgenic lines, facilitating a deeper understanding of parasite biology. The impact of genetic perturbations such as gene disruption or the introduction of drug resistance alleles on parasite fitness is typically quantified in competitive growth assays between the query line and a wild type reference. Although fluorescent reporter lines offer a facile and frequently used method to measure relative growth, this approach is limited by the strain background of the existing reporter, which may not match the growth characteristics of the query strains, particularly if these are slower-growing field isolates. Here, we demonstrate an efficient CRISPR-based approach to generate fluorescently labelled parasite lines using mNeonGreen derived from the LanYFP protein in *Branchiostoma lanceolatum*, which is one of the brightest monomeric green fluorescent proteins identified. Using a positive-selection approach by insertion of an in-frame blasticidin S deaminase marker, we generated a Dd2 reporter line expressing mNeonGreen under the control of the *pfpare* (*P. falciparum* Prodrug Activation and Resistance Esterase) locus. We selected the *pfpare* locus as an integration site because it is highly conserved across *P. falciparum* strains, expressed throughout the intraerythrocytic cycle, not essential, and offers the potential for negative selection to further enrich for integrants. The mNeonGreen@*pare* line demonstrates strong fluorescence with a negligible fitness defect. In addition, the construct developed can serve as a tool to fluorescently tag other *P. falciparum* strains for *in vitro* experimentation.

## 2 Introduction

Transgenic parasites expressing fluorescent proteins are a powerful tool in parasitology research. The ability to identify and track whole parasites or tagged parasite proteins *in vitro* has been integral for gaining insights into the biology of parasites and their interactions with hosts. In the study of malaria, the generation of fluorescent *Plasmodium* reporter lines has been instrumental in interrogating gene function, drug activity, life cycle, and host-parasite interactions (Portugaliza et al., 2019, Thommen et al., 2022, Talman et al., 2010, Frischknecht et al., 2006, Wilson et al., 2010, Voorberg-van der Wel et al., 2020). Reporter lines have also been instrumental in facilitating scaled-up analyses and the development of new methodologies. One important use of reporter lines is to quantify parasite fitness using competitive head-to-head growth assays. A query line, typically with an engineered mutation of interest such as a drug resistance allele or gene knockout, is mixed with a fluorescent isogenic wild type parasite. The change in relative abundance over time provides a measure of the fitness impact of the mutation of interest (Baragaña et al., 2015, Gabryszewski et al., 2016a, Ross et al., 2018, Stokes et al., 2021).

The first reporter lines developed in *Plasmodium* parasites expressed the exogenous proteins, firefly luciferase and chloramphenicol, using episomes (Goonewardene et al., 1993, Horrocks and Kilbey, 1996, Wu et al., 1995). Shortly after, green fluorescent protein (GFP) from *Aequorea victoria*, was developed as a reporter and adapted into *P. falciparum* and became widely used as it provided stable and strong fluorescence without requiring a cofactor (Chalfie et al., 1994, VanWye and Haldar, 1997). The reporter lines enabled functional and genetic analyses, particularly the bioluminescent and fluorescent reporters which supported imaging. The application of other reporters such as mCherry and yellow fluorescent protein and the transition to integrated reporters were gradual and hindered by challenges with homologous recombination-based integration (Engelmann et al., 2009, Armstrong and Goldberg, 2007). The advent of CRISPR/Cas9 genome editing and its application in *P. falciparum* made the development of reporter lines rapid and straightforward, allowing for the selective integration of fluorescent markers into specific genomic sites (Mogollon et al., 2016, Kuang et al., 2017, Miyazaki et al., 2020). Favourable integration sites could be selected based on the essentiality and phenotype of the gene and its expression profile, allowing for the development of reporter lines with stage-specific or multi-stage expression (Miyazaki et al., 2020, Marin-Mogollon et al., 2019). More recently developed fluorescent proteins i.e., mNeonGreen and mGreenLantern could generate better performing *Plasmodium* reporter lines as these proteins have a 3 and 6-fold increase in brightness compared to mEGFP, respectively. Faster maturation, improved acid tolerance, increased photostability and thermostability are also features that are often enhanced in these fluorescent proteins compared to standard fluorescent proteins (Shaner et al., 2013, Campbell et al., 2020).

In this work, we developed an efficient CRISPR/Cas9 approach to generate mNeonGreen-expressing *P. falciparum* lines, with the potential to use positive and negative selection to remove untagged parasites and obviate the need for clonal isolation. We inserted mNeonGreen in the genome at the non-essential *pfpare* (*P. falciparum* Prodrug Activation and Resistance Esterase) locus, so that its integration is stable, and its expression is endogenously driven by the *pfpare* promoter. We demonstrate that mNeonGreen@*pare* exhibits strong fluorescence throughout the intraerythrocytic cycle and has robust fitness compared to existing reporter lines.

## 3 Method

### 3.1 Sequence alignment

DNA sequences for *pfpare* locus (PF3D7_0709700) in *P. falciparum* 3D7, HB3, 7G8, GB4, CD01, GA01, IT and Dd2 strains were obtained from PlasmoDB and alignment figures were generated using Clustal Omega (Amos et al., 2021, Sievers et al., 2011, Goujon et al., 2010). Transcriptomic data of 3D7 *pfpare* expression in the intraerythrocytic life cycle was obtained from PlasmoDB, specifically RNA-sequencing data from Chappell *et al*. was used (Chappell et al., 2020).

### 3.2 Construct design and generation

The construct backbone used was pDC2-coCas9-gRNA, encoding Cas9, a gRNA expression cassette, hDHFR resistance for selection for plasmid uptake in *P. falciparum* culture and ampicillin resistance cassette for plasmid propagation in *E. coli* (Adjalley and Lee, 2022). Into this base vector, we inserted a guide RNA targeting *pfpare* (GGACAGTCAGAAGGATGGAA), and a donor region with flanking homology to 3D7 *pfpare* (255bp and 466bp homology regions; see Fig. 1B). We synthesised mNeonGreen codon-optimised for *P. falciparum* and cloned this downstream of the blasticidin S deaminase (BSD) selectable marker. To facilitate the expression of unfused proteins, 2A linkers were included separating the upstream *pfpare* fragment, BSD, and mNeonGreen (see Fig. 1B). Cloning was performed by Gibson assembly (NEBuilder DNA HiFi Assembly), and transformations were performed in XL-10 Gold Ultracompetent cells (Agilent). Plasmid sequences were confirmed by Sanger sequencing and plasmids were amplified and isolated using Midiprep plasmid extraction kits (Macherey-Nagel).

**Figure 1.**
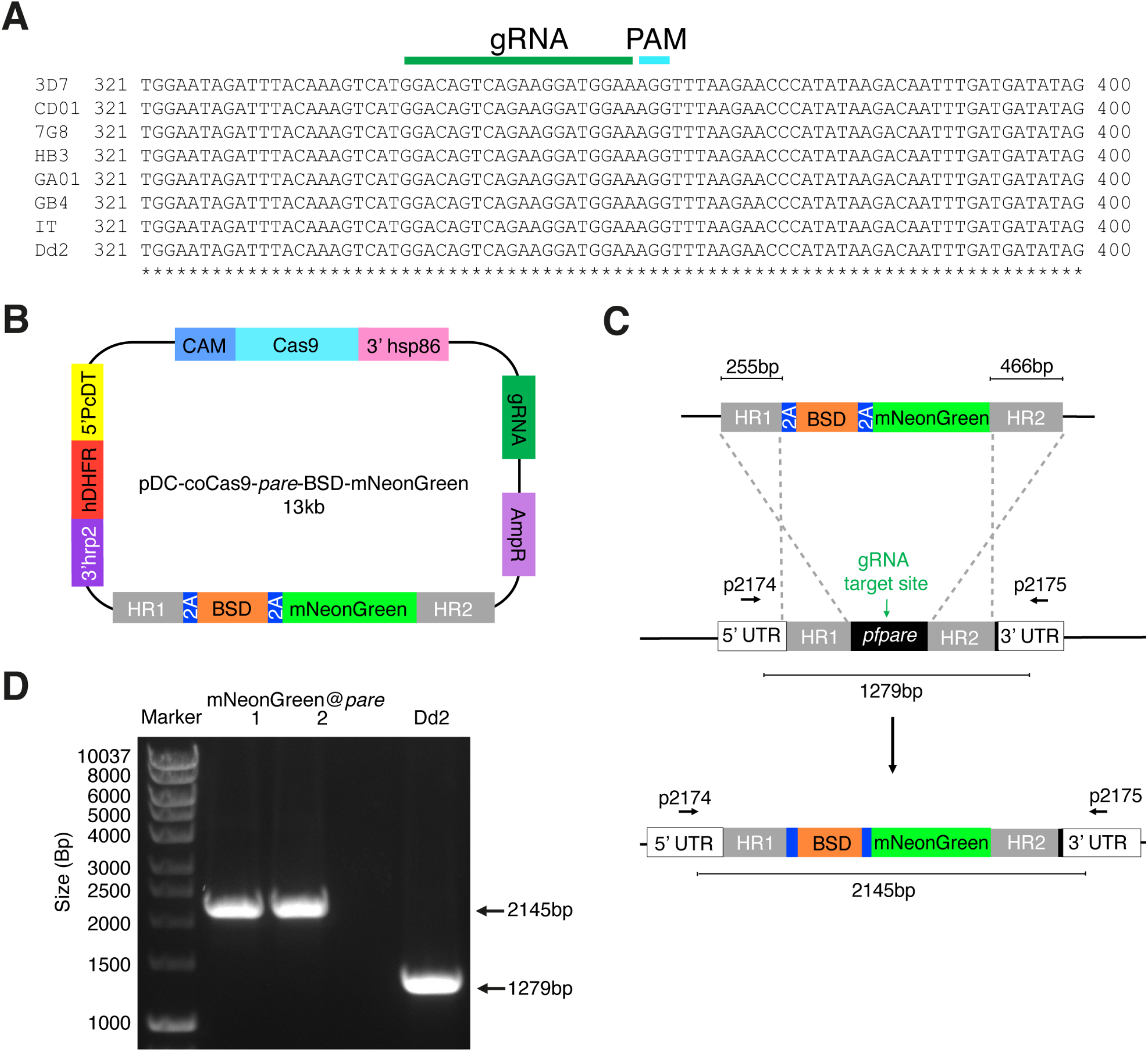
Generation of a *P. falciparum* Dd2 reporter line expressing mNeonGreen under the control of the highly conserved *pfpare* locus (mNeonGreen@*pare*). **(A)** DNA sequences for *pfpare* (PF3D7_0709700) in *P. falciparum* 3D7, HB3, 7G8, GB4, CD01, GA01, IT and Dd2 strains were obtained from PlasmoDB and Clustal Omega was used to create a multiple sequence alignment (Amos et al., 2021, Sievers et al., 2011, Goujon et al., 2010). **(B)** Schematic of the pDC2-coCas9-*pare*-BSD-mNeonGreen plasmid. The plasmid encodes a codon optimised Cas9 enzyme, guide RNA cassette containing a guide targeting *pfpare*, an ampicillin resistance cassette for plasmid propagation in *E. coli* and the hDHFR marker for selection for plasmid uptake in *P. falciparum*. The donor region contains a blasticidin S deaminase (BSD) selectable marker flanked by 2A linkers with mNeonGreen downstream. The payload is flanked by two homology regions for the *pfpare* locus. **(C)** Upon transfection of the pDC2-coCas9-*pare*-BSD-mNeonGreen plasmid into *P. falciparum* Dd2, the *pfpare*-targeting gRNA directs Cas9 to make a site-directed cut of *pfpare*. The donor region facilitates homology-directed repair and the insertion of BSD and mNeonGreen. **(D)** The *pfpare* locus of blasticidin-resistant recrudesced parasites (mNeonGreen@*pare*.1 and mNeonGreen@*pare*.2) was PCR-amplified using 5’ and 3’ UTR primers (p2174 and p2175) to check for correct insertion of mNeonGreen. Lengths of PCR products were quantified using gel electrophoresis. No detectable wild type product was observed in the transfected bulk culture.

### 3.3 Parasite culturing and plasmid transfections

Blood-stage *P. falciparum* parasites (Dd2, Dd2-EGFP and NF54-EGFP) were grown in RPMI media with AlbuMAX® (Gibco) and supplemented with GlutaMax® (Gibco), Gentamicin (Gibco) and HEPES (pH 7.0) with O+ human erythrocytes at 3% haematocrit. RBCs were obtained by anonymous donors from the National Health Services Blood and Transplant (NHSBT). Their use was in accordance with relevant guidelines and regulations, with approval from the NHS Cambridgeshire Research Ethics Committee and the Wellcome Sanger Institute Human Materials and Data Management Committee. Cultures were maintained in a gaseous environment of 3% CO_2_, 1% O_2_ and 96% N_2_ at 37°C. Parasitemia and stages were monitored using Giemsa staining and microscopy. Synchronisation was completed using sorbitol ring enrichment (Radfar et al., 2009). Transfections were completed by electroporation of parasitised red blood cells (Bio-Rad Gene Pulser Xcell) as previously described (Fidock and Wellems, 1997). Cultures containing 5% ring-stage parasites were transfected with 50µg of the plasmid. Blasticidin S drug selection (2µg/ml) was applied one-day post-electroporation and maintained continuously. Correct editing of recrudesced parasites (mNeonGreen@*pare*.1 and mNeonGreen@*pare*.2) was confirmed using primers (p2174 and p2175), which flank the 5’ and 3’ UTRs of *pfpare* (TGCACTTGTTTTACATTTTTATATT and TGTAACATCACTAATTAATTTATTAA). PCR amplification and gel electrophoresis were used to check insert size and Sanger sequencing to confirm the correct sequence. The Dd2-EGFP and NF54-EGFP fluorescent lines were previously published (Baragaña et al., 2015, Gabryszewski et al., 2016b). Both lines were generated using attB x attP recombination to insert the reporter genes, however, they are driven by different constitutive promoters: Dd2-EGFP is driven by the ER hsp70 (PF3D7_0917900) promoter and the NF54-EGFP by the calmodulin (PF3D7_1434200) promoter.

### 3.4 Fluorescence microscopy

200µL of parasite culture was harvested at 5% parasitemia, spun at 3000rpm for 30 seconds in an Eppendorf tube and washed with 0.5mL of PBS. Parasites were fixed by resuspending in 4% (v/v) paraformaldehyde + 0.0075% glutaraldehyde, incubating for 30 minutes, and washed twice with 1mL of PBS. Parasite DNA was stained with 1mL of Hoechst stain (10µg/mL) for 5 minutes and resuspended in 1mL of PBS. The stained parasites were transferred to an 8-well chambered coverglass (Lab-Tek) pre-treated with 0.25mL of poly L-lysine (0.2mg/ml) for 10mins. The coverglass was transferred to an inverted fluorescence microscope (Leica DMi8). Bright-field microscopy and blue and green-filtered fluorescence microscopy were used with the 100x objective (total 1000x magnification) to image infected erythrocytes and fluorescent parasites. Image processing and analysis were completed with Leica Application Suite (LAS X).

### 3.5 Flow cytometric analysis of fluorescence

Parasites were transferred to a round-bottom 96-well plate, spun (3min s, 2000rpm) and the supernatant was removed. To check viability, infected erythrocytes were stained with 100nM MitoTracker Deep Red (MitoDR) solution (Thermofisher) in NaCl 0.9%/Dextrose 0.2% and incubated for 30mins in the dark at 37°C. After a 1/40 dilution in PBS, parasites were analysed on a flow cytometer (Beckman Coulter CytoFLEX S) and at least 20 000 events were recorded for each experiment. The blue 488nm laser was used to detect green fluorescence and the red 638nm laser was used to detect MitoDR. Quantification of green fluorescent (GFP and mNeonGreen) and MitoDR positive cells in the culture population was performed using FlowJo (v10).

### 3.6 Competition assay

Pre-assay parasitemia was measured using flow cytometry and stages were observed with microscopy. Mixed-stage lines of mNeonGreen@*pare*.1, mNeonGreen@*pare*.2 and Dd2-EGFP were competed against Dd2 or 3D7 in triplicate at 50:50 starting ratio with a total of 1% parasitemia. The parasitemia and fluorescence were measured using flow cytometry every 2^nd^ or 3^rd^ day from day 0 to day 21. Parasitemia was maintained between 0.5%-6%. The percentage of parasites expressing green fluorescence over total parasites was averaged between the triplicates and graphed over time for each line.

## 4 Results

### 4.1 Selection of the *pfpare* locus for fluorescent markers integration in *P. falciparum*

The *pfpare* locus was identified as an optimal safe-harbour site for the integration of the mNeonGreen fluorescent marker. *pfpare* (PF3D7_0709700) encodes a prodrug activation and resistance esterase that is not essential (Istvan et al., 2017). In addition to the dispensible nature of *pfpare*, loss-of-function mutations in *pfpare* confer resistance to the antimalarial compound MMV011438, which requires *pfpare* for its activation. *pfpare* is expressed during blood stages, therefore its promoter would facilitate the expression of a fluorescent marker throughout the intraerythrocytic life cycle. In addition, we included in the inserted sequence the blasticidin S deaminase (BSD) marker, which would only be expressed from the *pfpare* promoter once integrated. Thus our strategy would permit both positive and negative selection options if required for efficient isolation of a homogenous culture of mNeonGreen tagged parasites without cloning.

To identify if *pfpare* is suitable as an integration site across multiple *P. falciparum* strains, a multiple sequence alignment of the *pfpare* locus for eight geographically diverse strains (3D7, HB3, 7G8, GB4, CD01, GA01, IT and Dd2) was completed. We first identified a gRNA target site that was highly conserved, with no mutations in the guide RNA or PAM sequences (Fig. 1A). Examination of the flanking upstream and downstream sequences that would constitute the donor homology regions revealed no mutations in the 5’ homology region and 2 − 3 single nucleotide polymorphisms in the 3’ homology region of the donor sequence of the plasmid (Supplemental Fig. 1). This level of sequence diversity would not be expected to strongly impact editing. We validated this prediction below using a 3D7-based donor sequence to edit Dd2, which has three polymorphisms relative to 3D7.

### 4.2 Generation of an endogenous mNeonGreen-expressing *P. falciparum* reporter line

To design a construct to integrate mNeonGreen into *pfpare*, we first subcloned a codon-optimised mNeonGreen downstream of BSD, with flanking 5’ and 3’ *pfpare* (3D7) homology regions of 255 bp and 466 bp respectively. The resulting plasmid (pDC-coCas9-*pare*-BSD-mNeonGreen; Fig. 1B) expresses Cas9 driven by the calmodulin promoter and transcribes a *pfpare*-targeting guide RNA. Plasmids were transfected into *P. falciparum* Dd2 parasites and continuous selection with blasticidin S was used to select for plasmid uptake and the insertion of the donor into *pfpare* by homology-directed repair (Fig. 1C). Edited parasites were obtained from two different transfections, referred to as mNeonGreen@*pare*.1 and mNeonGreen@*pare*.2. PCR-amplification of the *pfpare* locus demonstrated successful integration of the donor with no wild type locus detectable, indicating positive selection was sufficient to deplete any unedited parasites (Fig. 1D, primers shown in Fig. 1C).

### 4.3 Endogenous mNeonGreen expression generates fluorescence comparable to GFP

To determine if the integrated mNeonGreen yields fluorescent parasites, we first examined the transgenic lines by fluorescence microscopy. Infected erythrocytes were detected using Hoechst DNA stain, which does not stain uninfected erythrocytes because they are anucleate, unlike *Plasmodium* parasites. Erythrocytes infected with either of the mNeonGreen@*pare* lines showed strong green fluorescence unlike the parental Dd2 line (Fig. 2A). Flow cytometry was used to quantify the level of fluorescence and distribution within the population of a mixed-stage culture. The fluorescence profiles of mNeonGreen@*pare*.1 and mNeonGreen@*pare*.2 were highly similar and are comparable to the profiles of other green-fluorescing lines used for competition assays, including Dd2-EGFP and NF54-EGFP, which express GFP from the strong constitutive promoters of ER-Hsp70 and calmodulin respectively (Adjalley et al., 2010, Baragaña et al., 2015). The peaks of the mNeonGreen@*pare* lines were modestly shifted left in comparison to the GFP lines, which means that the bulk of mNeonGreen@*pare* parasites in mixed culture are less fluorescent than the bulk of GFP-expressing parasites but still readily distinguishable from non-fluorescent parasites (Fig. 2B). The bimodal peaks suggested that subpopulations expressed different levels of fluorescence.

**Figure 2.**
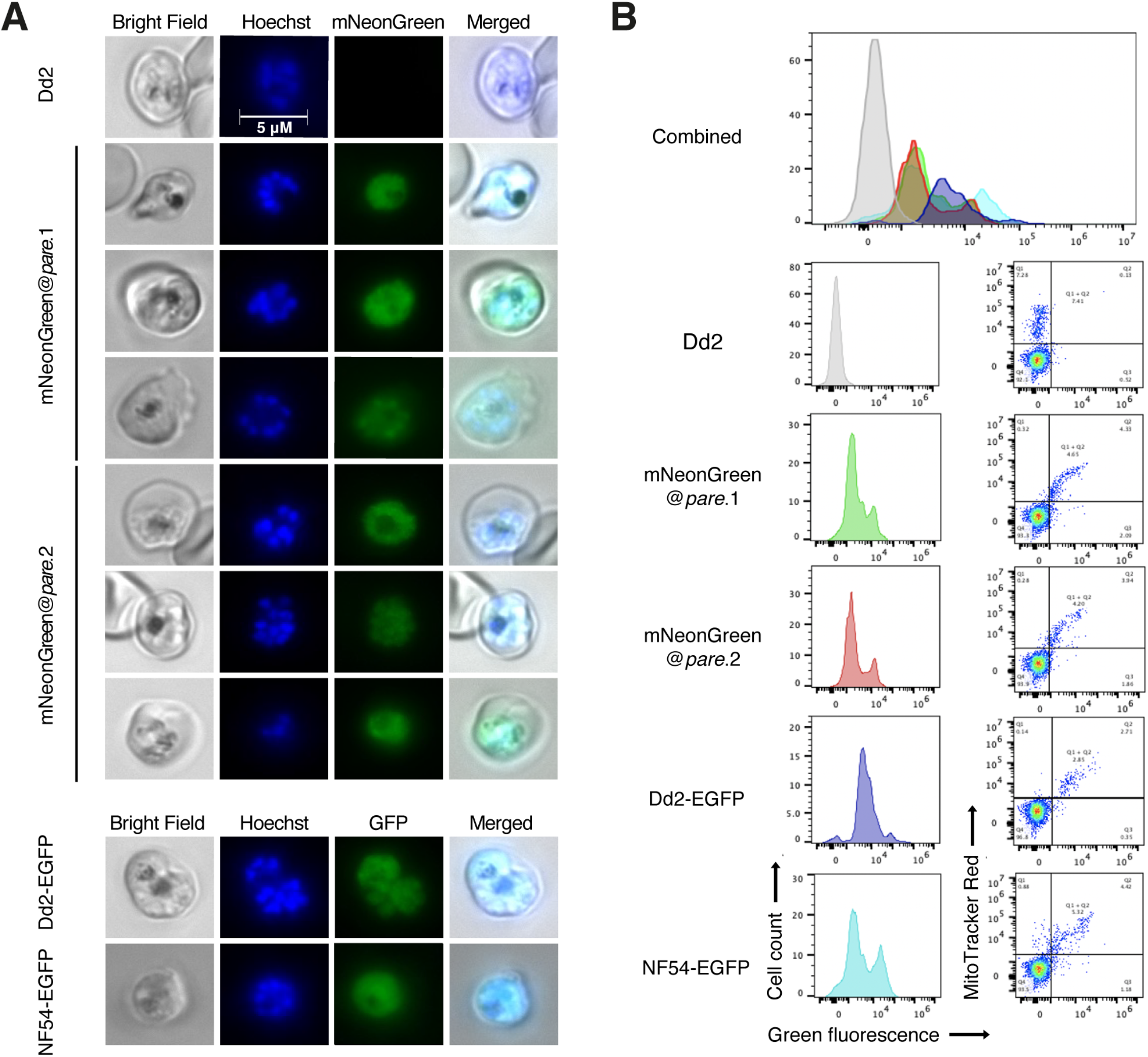
mNeonGreen fluorescence in *P. falciparum* is comparable to GFP fluorescent lines. **(A)** Mixed-staged parasites (mNeonGreen@*pare*.1, mNeonGreen@*pare*.2, Dd2, NF54-EGFP, Dd2-EGFP) were fixed using 4% (v/v) paraformaldehyde + 0.0075% glutaraldehyde, stained with Hoechst stain, and placed on a poly L-lysine coated coverglass for inverted fluorescence microscopy. Bright-field microscopy and fluorescence microscopy were used with a 100x objective (total 1000x magnification) and with green and blue fluorescence was captured to image infected RBCs and fluorescent parasites. **(B)** Mixed-stage parasites (mNeonGreen@pare.1, mNeonGreen@pare.2, Dd2, NF54-EGFP, Dd2-EGFP) were stained with MitoTracker DeepRed and analysed on a flow cytometer. Quantification of green fluorescent (GFP and mNeonGreen) and MitoTracker^+^ cells was performed using FlowJo. All single-cell, parasitised RBCs (MitoTracker^+^) were gated, and histograms of green fluorescence were generated (GFP^+^ or mNeonGreen^+^).

### 4.4 mNeonGreen fluorescence varies between different asexual *P. falciparum* blood stages

In a healthy asexual intraerythrocytic *P. falciparum* culture, the subpopulations include briefly free-roaming merozoites and intraerythrocytic ring, trophozoite and schizont stages. To assess the level of fluorescence in specific stages, we enriched mNeonGreen@*pare* cultures for different stages using synchronisation. Stage-specific levels of fluorescence were observed with schizonts producing the highest fluorescence, followed by trophozoites and lastly, ring stages (Fig. 3A). This pattern was consistent with transcriptomic studies that demonstrate that the expression of *pfpare* peaks at 32-40 hours post-erythrocyte-invasion, which would suggest that the expression of mNeonGreen from the *pfpare* locus would also peak during late trophozoite and schizont stages (Fig 3B) (Chappell et al., 2020, Amos et al., 2021).

**Figure 3.**
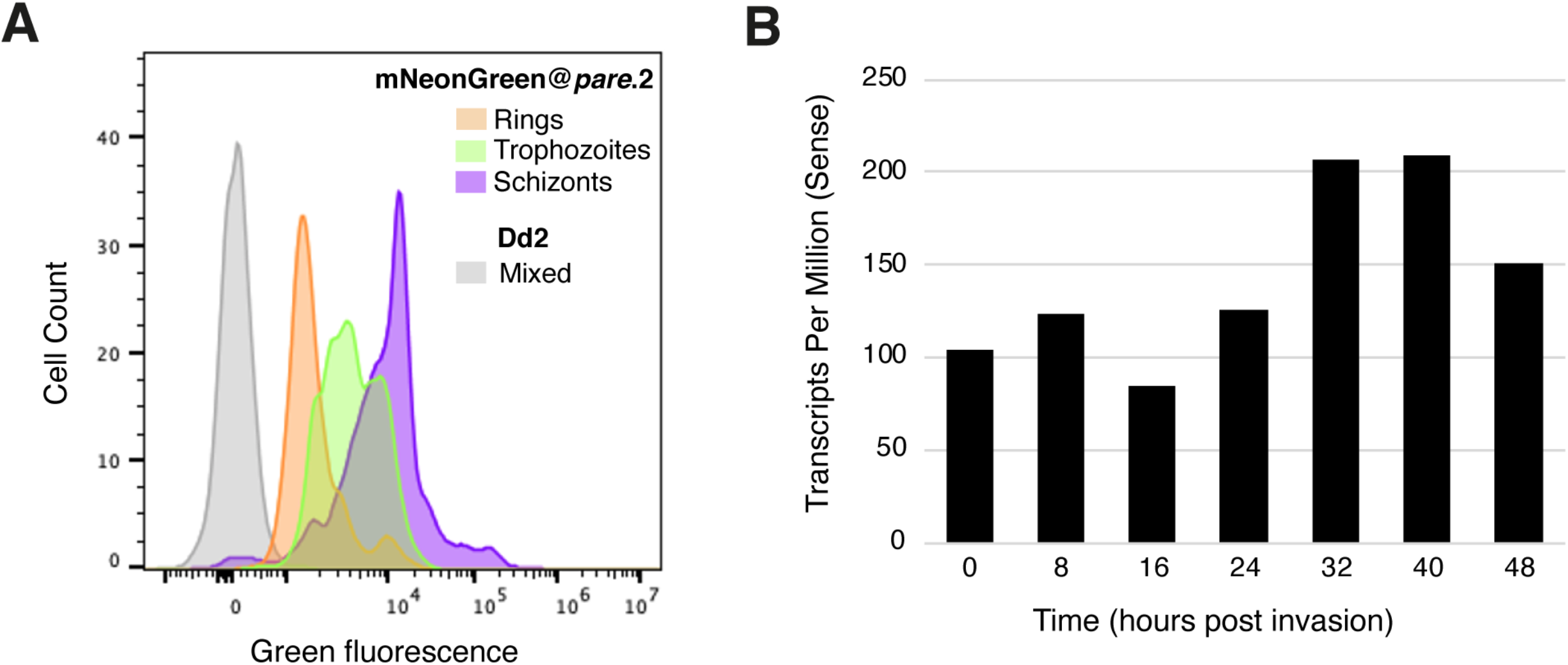
mNeonGreen@*pare* fluorescence varies between different asexual blood stages. (**A)** mNeonGreen@*pare*.2 parasites were synchronised using sorbitol ring-stage enrichment and microscopy was used to confirm stages. Flow cytometry with MitoTracker DeepRed was used to enumerate parasites expressing green fluorescence in ring, trophozoite and schizont-staged cultures. FlowJo was used to gate single-cell, parasitised RBCs, quantify green fluorescence, and generate histograms. mNeonGreen@*pare* schizonts demonstrated the greatest fluorescence followed by trophozoites and then rings. (**B)** Stage-specific RNA-sequencing obtained from PlasmoDB shows that *pfpare* is expressed throughout the entire 48hr intraerythrocytic life cycle and expression peaks at late trophozoite and schizont stages (Amos et al., 2021, Chappell et al., 2020).

### 4.5 mNeonGreen@*pare* lines demonstrate robust fitness

To assess the fitness of the mNeonGreen@*pare* lines, competition assays were performed against Dd2 and 3D7. The two populations were seeded at a 1:1 ratio of fluorescent to test line and maintained for 3 weeks. The ratio of fluorescent (mNeonGreen@*pare*) to non-fluorescent (Dd2 or 3D7) populations was measured every 2 - 3 days using flow cytometry to quantify the competition between the two populations. Dd2-EGFP was also included as a control, which has been shown previously to have a slight fitness defect due to the integration of GFP (Baragaña et al., 2015). The mNeonGreen@*pare* lines showed nearly comparable fitness to their parent line, Dd2, over a three-week period (Fig. 4A). Contrastingly when competed against 3D7, a slower-growing lab line, the mNeonGreen@*pare* lines outcompeted 3D7, which demonstrates the value of strain-matched competitor lines (Fig. 4B).

**Figure 4.**
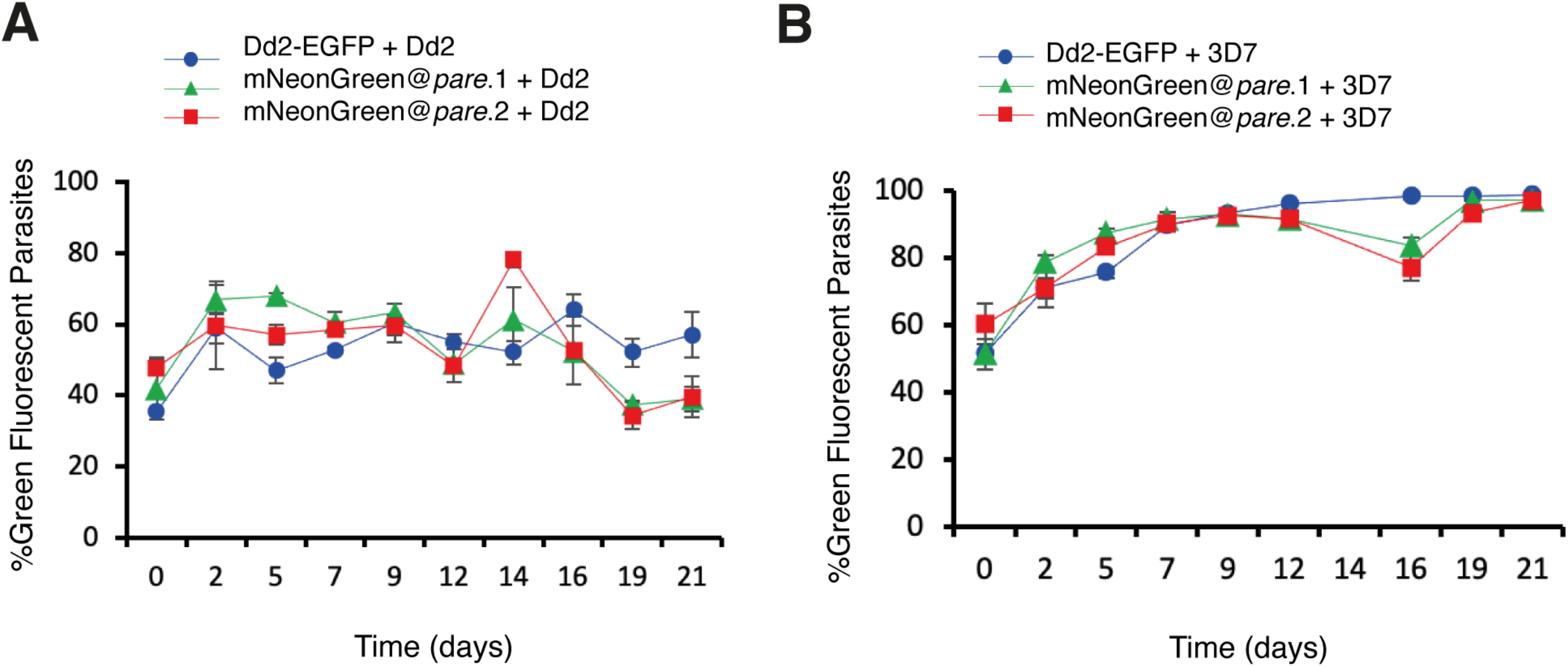
mNeonGreen@*pare* lines demonstrate comparable fitness to Dd2-EGFP when competed against Dd2 or 3D7. Fluorescent lines (mNeonGreen@pare.1, mNeonGreen@pare.2 and Dd2-EGFP) were competed against (**A**) Dd2 or (**B**) 3D7 in triplicate. The two populations were seeded at a 1:1 ratio at 1% parasitemia and maintained for 3 weeks. The parasitemia and fluorescence were measured using flow cytometry every 2^nd^ or 3^rd^ day from day 0 to day 21. The percentage of green fluorescent parasites over total parasites was averaged between the triplicates and graphed over time for each line. In the competition assay vs. Dd2, the three fluorescent lines did not outcompete Dd2 and remained close to the seeded percentages over three weeks. However, in the competition assay vs. 3D7, the three fluorescent lines quickly outcompeted 3D7.

## 5 Discussion

Genetic engineering of fluorescent proteins has greatly improved their functionality as tools in molecular biology. As such the integration of these enhanced proteins into existing applications in malaria research should be explored. In this work, we have developed an efficient and facile approach for generating new fluorescent reporter lines in *P. falciparum* using CRISPR-based integration of mNeonGreen. mNeonGreen was first applied to *P. falciparum* to generate a tagged Exp2 protein however, here we aimed to assess its broader applicability in generating reporter lines for competitive fitness assays (Glushakova et al., 2018). Therefore, we designed a CRISPR/Cas9 construct to integrate mNeonGreen into *pfpare*, a highly conserved, nonessential gene that would endogenously drive expression throughout the intraerythrocytic cycle.

We used the construct to tag the Dd2 strain and then assessed mNeonGreen@*pare* for features suitable in a reporter line i.e., the strength of fluorescence, localisation, stage-specificity, and impact on parasite fitness. mNeonGreen@*pare* demonstrated strong fluorescence that was diffused throughout the parasite. The fluorescence was expressed in all asexual blood stages but was most concentrated in schizonts. The ability to efficiently tag Dd2 indicates that minor differences in the homology regions between the donor, based on the 3D7 sequence, and the target region are tolerated. The absence of wild type locus in bulk transfections reflects the effect of positive selection resulting from *bsd* expression from the endogenous *pfpare* promoter. Although disruption of *pfpare* also affords the possibility of negative selection using the commercially available compound MMV011438, this was not required in practice due to the stringency of the blasticidin S positive selection. The integration of mNeonGreen into *pfpare* caused a minor fitness defect leading to slightly slower growth compared to Dd2, similar to other GFP-based reporters (Baragaña et al., 2015). However, this defect was relatively minimal, as the resulting line outcompeted 3D7 in a competition assay. These findings support that the mNeonGreen@*pare* is a suitable reporter line and is comparable with the standard *P. falciparum* GFP lines that are currently used.

Both tools generated in this work, the Dd2 mNeonGreen reporter line and the construct that generates new reporter lines, will support malaria research. mNeonGreen@*pare* is a valuable addition to the repertoire of reporter lines in *P. falciparum* that facilitate experiments involving visualising, tracking and counting parasites. The pDC-coCas9-*pare*-BSD-mNeonGreen construct will enable the rapid generation of other fluorescently tagged *P. falciparum* parasites from different strains, such as field isolates, which can facilitate their study *in vitro*.

## 7 Author Contributions

JH and ML conceived and designed the experiments. HJ generated the pDC2-coCas9-*pare*-2A-BSD construct. JH generated the pDC-coCas9-*pare*-BSD-mNeonGreen plasmid and performed the experiments and analysis. ML supervised the work. JH and ML wrote the manuscript. All authors read and approved the manuscript.

## 10 Figure Legends

**Supplemental Figure 1.**
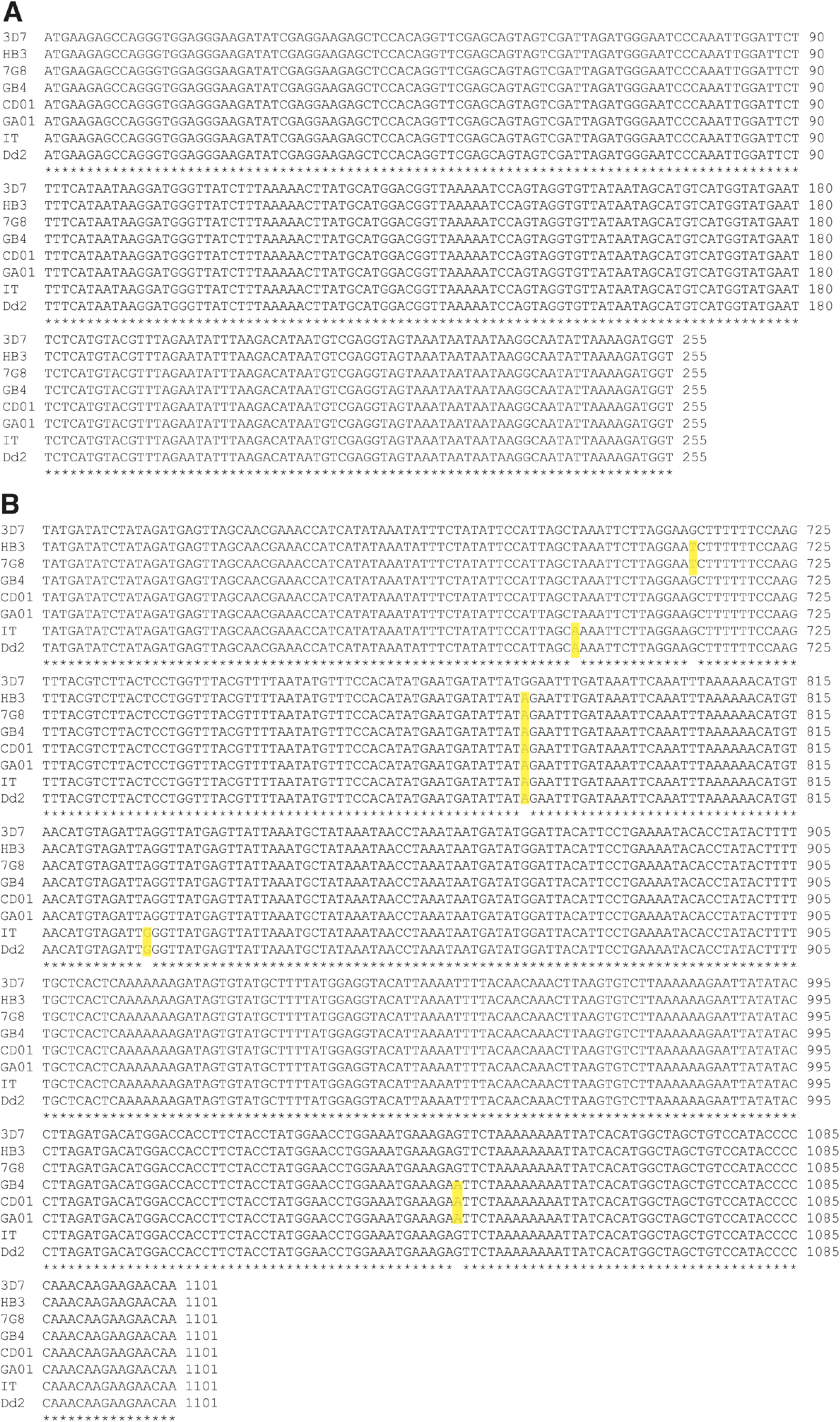
Alignment of 5’ and 3’ homologous regions of the *pfpare* locus in *P. falciparum* sequences. The DNA sequences of the 5’ (**A**) and 3’ (**B**) *pfpare* homologous regions used in pDC2-coCas9-*pare*-BSD-mNeonGreen were obtained for *P. falciparum* 3D7, HB3, 7G8, GB4, CD01, GA01, IT and Dd2 strains from PlasmoDB (Amos et al., 2021). Clustal Omega was used to create a multiple sequence alignment (Sievers et al., 2011, Goujon et al., 2010). Single nucleotide polymorphisms are highlighted with yellow boxes.

## References

Adjalley, S. H. & Lee, M. C. 2022. CRISPR/Cas9 editing of the Plasmodium falciparum genome. Methods in Molecular Biology.

Adjalley, S. H., Lee, M. C. & Fidock, D. A. 2010. A method for rapid genetic integration into Plasmodium falciparum utilizing mycobacteriophage Bxb1 integrase. Methods Mol Biol, 634, 87–100.

Amos, B., Aurrecoechea, C., Barba, M., Barreto, A., Basenko Evelina y., Bazant, W., Belnap, R., Blevins, A. S., Böhme, U., Brestelli, J., Brunk, B. P., Caddick, M., Callan, D., Campbell, L., Christensen Mikkel b., Christophides George k., Crouch, K., Davis, K., Debarry, J., Doherty, R., Duan, Y., Dunn, M., Falke, D., Fisher, S., Flicek, P., Fox, B., Gajria, B., Giraldo-Calderón, G. I., Harb, O. S., Harper, E., Hertz-Fowler, C., Hickman Mark j., Howington, C., Hu, S., Humphrey, J., Iodice, J., Jones, A., Judkins, J., Kelly, S. A., Kissinger, J. C., Kwon, D. K., Lamoureux, K., Lawson, D., Li, W., Lies, K., Lodha, D., Long, J., Maccallum, R. M., Maslen, G., Mcdowell, M. A., Nabrzyski, J., Roos, D. S., Rund, S. S. C., Schulman Stephanie w., Shanmugasundram, A., Sitnik, V., Spruill, D., Starns, D., Stoeckert Christian j., Jr, Tomko, S. S., Wang, H., Warrenfeltz, S., Wieck, R., Wilkinson, P. A., Xu, L. & Zheng, J. 2021. VEuPathDB: the eukaryotic pathogen, vector and host bioinformatics resource center. Nucleic Acids Research, 50, D898–D911.

Armstrong, C. M. & Goldberg, D. E. 2007. An FKBP destabilization domain modulates protein levels in Plasmodium falciparum. Nat Methods, 4, 1007–9.

Baragaña, B., Hallyburton, I., Lee, M. C., Norcross, N. R., Grimaldi, R., Otto, T. D., Proto, W. R., Blagborough, A. M., Meister, S., Wirjanata, G., Ruecker, A., Upton, L. M., Abraham, T. S., Almeida, M. J., Pradhan, A., Porzelle, A., Luksch, T., Martínez, M. S., Luksch, T., Bolscher, J. M., Woodland, A., Norval, S., Zuccotto, F., Thomas, J., Simeons, F., Stojanovski, L., Osuna-Cabello, M., Brock, P. M., Churcher, T. S., Sala, K. A., Zakutansky, S. E., Jiménez-Díaz, M. B., Sanz, L. M., Riley, J., Basak, R., Campbell, M., Avery, V. M., Sauerwein, R. W., Dechering, K. J., Noviyanti, R., Campo, B., Frearson, J. A., Angulo-Barturen, I., Ferrer-Bazaga, S., Gamo, F. J., Wyatt, P. G., Leroy, D., Siegl, P., Delves, M. J., Kyle, D. E., Wittlin, S., Marfurt, J., Price, R. N., Sinden, R. E., Winzeler, E. A., Charman, S. A., Bebrevska, L., Gray, D. W., Campbell, S., Fairlamb, A. H., Willis, P. A., Rayner, J. C., Fidock, D. A., Read, K. D. & Gilbert, I. H. 2015. A novel multiple-stage antimalarial agent that inhibits protein synthesis. Nature, 522, 315–20.

Campbell, B. C., Nabel, E. M., Murdock, M. H., Lao-Peregrin, C., Tsoulfas, P., Blackmore, M. G., Lee, F. S., Liston, C., Morishita, H. & Petsko, G. A. 2020. mGreenLantern: a bright monomeric fluorescent protein with rapid expression and cell filling properties for neuronal imaging. Proc Natl Acad Sci U S A, 117, 30710–30721.

Chalfie, M., Tu, Y., Euskirchen, G., Ward, W. W. & Prasher, D. C. 1994. Green Fluorescent Protein as a Marker for Gene Expression. Science, 263, 802–805.

Chappell, L., Ross, P., Orchard, L., Russell, T. J., Otto, T. D., Berriman, M., Rayner, J. C. & Llinás, M. 2020. Refining the transcriptome of the human malaria parasite Plasmodium falciparum using amplification-free RNA-seq. BMC Genomics, 21, 395.

Engelmann, S., Silvie, O. & Matuschewski, K. 2009. Disruption of Plasmodium sporozoite transmission by depletion of sporozoite invasion-associated protein 1. Eukaryot Cell, 8, 640–8.

Fidock, D. A. & Wellems, T. E. 1997. Transformation with human dihydrofolate reductase renders malaria parasites insensitive to WR99210 but does not affect the intrinsic activity of proguanil. Proc Natl Acad Sci U S A, 94, 10931–6.

Frischknecht, F., Martin, B., Thiery, I., Bourgouin, C. & Menard, R. 2006. Using green fluorescent malaria parasites to screen for permissive vector mosquitoes. Malaria Journal, 5, 23.

Gabryszewski, S. J., Dhingra, S. K., Combrinck, J. M., Lewis, I. A., Callaghan, P. S., Hassett, M. R., Siriwardana, A., Henrich, P. P., Lee, A. H., Gnädig, N. F., Musset, L., Llinás, M., Egan, T. J., Roepe, P. D. & Fidock, D. A. 2016a. Evolution of Fitness Cost-Neutral Mutant PfCRT Conferring P. falciparum 4-Aminoquinoline Drug Resistance Is Accompanied by Altered Parasite Metabolism and Digestive Vacuole Physiology. PLoS Pathog, 12, e1005976.

Gabryszewski, S. J., Modchang, C., Musset, L., Chookajorn, T. & Fidock, D. A. 2016b. Combinatorial Genetic Modeling of pfcrt-Mediated Drug Resistance Evolution in Plasmodium falciparum. Molecular Biology and Evolution, 33, 1554–1570.

Glushakova, S., Beck, J. R., Garten, M., Busse, B. L., Nasamu, A. S., Tenkova-Heuser, T., Heuser, J., Goldberg, D. E. & Zimmerberg, J. 2018. Rounding precedes rupture and breakdown of vacuolar membranes minutes before malaria parasite egress from erythrocytes. Cell Microbiol, 20, e12868.

Goonewardene, R., Daily, J., Kaslow, D., Sullivan, T. J., Duffy, P., Carter, R., Mendis, K. & Wirth, D. 1993. Transfection of the malaria parasite and expression of firefly luciferase. Proc Natl Acad Sci U S A, 90, 5234–6.

Goujon, M., Mcwilliam, H., Li, W., Valentin, F., Squizzato, S., Paern, J. & Lopez, R. 2010. A new bioinformatics analysis tools framework at EMBL-EBI. Nucleic Acids Research, 38, W695–W699.

Horrocks, P. & Kilbey, B. J. 1996. Physical and functional mapping of the transcriptional start sites of Plasmodium falciparum proliferating cell nuclear antigen. Mol Biochem Parasitol, 82, 207–15.

Istvan, E. S., Mallari, J. P., Corey, V. C., Dharia, N. V., Marshall, G. R., Winzeler, E. A. & Goldberg, D. E. 2017. Esterase mutation is a mechanism of resistance to antimalarial compounds. Nature Communications, 8, 14240.

Kuang, D., Qiao, J., Li, Z., Wang, W., Xia, H., Jiang, L., Dai, J., Fang, Q. & Dai, X. 2017. Tagging to endogenous genes of Plasmodium falciparum using CRISPR/Cas9. Parasit Vectors, 10, 595.

Marin-Mogollon, C., Salman, A. M., Koolen, K. M. J., Bolscher, J. M., Van Pul, F. J. A., Miyazaki, S., Imai, T., Othman, A. S., Ramesar, J., Van Gemert, G. J., Kroeze, H., Chevalley-Maurel, S., Franke-Fayard, B., Sauerwein, R. W., Hill, A. V. S., Dechering, K. J., Janse, C. J. & Khan, S. M. 2019. A P. falciparum NF54 Reporter Line Expressing mCherry-Luciferase in Gametocytes, Sporozoites, and Liver-Stages. Front Cell Infect Microbiol, 9, 96.

Miyazaki, S., Yang, A. S. P., Geurten, F. J. A., Marin-Mogollon, C., Miyazaki, Y., Imai, T., Kolli, S. K., Ramesar, J., Chevalley-Maurel, S., Salman, A. M., Van Gemert, G. A., Van Waardenburg, Y. M., Franke-Fayard, B., Hill, A. V. S., Sauerwein, R. W., Janse, C. J. & Khan, S. M. 2020. Generation of Novel Plasmodium falciparum NF135 and NF54 Lines Expressing Fluorescent Reporter Proteins Under the Control of Strong and Constitutive Promoters. Front Cell Infect Microbiol, 10, 270.

Mogollon, C. M., Van Pul, F. J., Imai, T., Ramesar, J., Chevalley-Maurel, S., De Roo, G. M., Veld, S. A., Kroeze, H., Franke-Fayard, B. M., Janse, C. J. & Khan, S. M. 2016. Rapid Generation of Marker-Free P. falciparum Fluorescent Reporter Lines Using Modified CRISPR/Cas9 Constructs and Selection Protocol. PLoS One, 11, e0168362.

Portugaliza, H. P., Llorà-Batlle, O., Rosanas-Urgell, A. & Cortés, A. 2019. Reporter lines based on the gexp02 promoter enable early quantification of sexual conversion rates in the malaria parasite Plasmodium falciparum. Scientific Reports, 9, 14595.

Radfar, A., Méndez, D., Moneriz, C., Linares, M., Marín-García, P., Puyet, A., Diez, A. & Bautista, J. M. 2009. Synchronous culture of Plasmodium falciparum at high parasitemia levels. Nature Protocols, 4, 1899–1915.

Ross, L. S., Dhingra, S. K., Mok, S., Yeo, T., Wicht, K. J., Kümpornsin, K., Takala-Harrison, S., Witkowski, B., Fairhurst, R. M., Ariey, F., Menard, D. & Fidock, D. A. 2018. Emerging Southeast Asian PfCRT mutations confer Plasmodium falciparum resistance to the first-line antimalarial piperaquine. Nat Commun, 9, 3314.

Shaner, N. C., Lambert, G. G., Chammas, A., Ni, Y., Cranfill, P. J., Baird, M. A., Sell, B. R., Allen, J. R., Day, R. N., Israelsson, M., Davidson, M. W. & Wang, J. 2013. A bright monomeric green fluorescent protein derived from Branchiostoma lanceolatum. Nat Methods, 10, 407–9.

Sievers, F., Wilm, A., Dineen, D., Gibson, T. J., Karplus, K., Li, W., Lopez, R., Mcwilliam, H., Remmert, M., Söding, J., Thompson, J. D. & Higgins, D. G. 2011. Fast, scalable generation of high-quality protein multiple sequence alignments using Clustal Omega. Molecular Systems Biology, 7, 539.

Stokes, B. H., Dhingra, S. K., Rubiano, K., Mok, S., Straimer, J., Gnädig, N. F., Deni, I., Schindler, K. A., Bath, J. R., Ward, K. E., Striepen, J., Yeo, T., Ross, L. S., Legrand, E., Ariey, F., Cunningham, C. H., Souleymane, I. M., Gansané, A., Nzoumbou-Boko, R., Ndayikunda, C., Kabanywanyi, A. M., Uwimana, A., Smith, S. J., Kolley, O., Ndounga, M., Warsame, M., Leang, R., Nosten, F., Anderson, T. J., Rosenthal, P. J., Ménard, D. & Fidock, D. A. 2021. Plasmodium falciparum K13 mutations in Africa and Asia impact artemisinin resistance and parasite fitness. Elife, 10.

Talman, A. M., Blagborough, A. M. & Sinden, R. E. 2010. A Plasmodium falciparum strain expressing GFP throughout the parasite’s life-cycle. PLoS One, 5, e9156.

Thommen, B. T., Passecker, A., Buser, T., Hitz, E., Voss, T. S. & Brancucci, N. M. B. 2022. Revisiting the Effect of Pharmaceuticals on Transmission Stage Formation in the Malaria Parasite Plasmodium falciparum. Frontiers in Cellular and Infection Microbiology, 12.

Vanwye, J. D. & Haldar, K. 1997. Expression of green fluorescent protein in Plasmodium falciparum. Mol Biochem Parasitol, 87, 225–9.

Voorberg-Van Der Wel, A. M., Zeeman, A.-M., Nieuwenhuis, I. G., Van Der Werff, N. M., Klooster, E. J., Klop, O., Vermaat, L. C., Kumar Gupta, D., Dembele, L., Diagana, T. T. & Kocken, C. H. M. 2020. A dual fluorescent Plasmodium cynomolgi reporter line reveals in vitro malaria hypnozoite reactivation. Communications Biology, 3, 7.

Wilson, D. W., Crabb, B. S. & Beeson, J. G. 2010. Development of fluorescent Plasmodium falciparum for in vitro growth inhibition assays. Malaria Journal, 9, 152.

Wu, Y., Sifri, C. D., Lei, H. H., Su, X. Z. & Wellems, T. E. 1995. Transfection of Plasmodium falciparum within human red blood cells. Proc Natl Acad Sci U S A, 92, 973–7.

